# Integrative single-cell and bulk RNA-seq analysis in human retina identified cell type-specific composition and gene expression changes for age-related macular degeneration

**DOI:** 10.1101/768143

**Authors:** Yafei Lyu, Randy Zauhar, Nico Dana, Christianne E. Strang, Kui Wang, Shanrun Liu, Zhen Miao, Naifei Pan, Paul Gamlin, James A. Kimble, Jeffrey D. Messinger, Christine A. Curcio, Dwight Stambolian, Mingyao Li

**Author notes:** Equal contribution (Yafei Lyu, Randy Zauhar).

## Abstract

Age-related macular degeneration (AMD) preferentially affects distinct cell types and topographic regions in retina. To characterize the impact of AMD on gene expression changes across retinal cell types and regions, we generated both single-cell RNA-seq (scRNA-seq) and bulk RNA-seq data from macular and peripheral retina in postmortem human donors with and without AMD. The scRNA-seq data revealed 11 major cell types with many previously reported AMD risk genes showing substantial cell type and region specificity. Cell type proportional changes with advancing AMD stage were significant for Müller glia, rods, astrocytes, microglia and endothelium.

---

AMD affects over 10 million Americans^1^, twice the number affected by Alzheimer’s disease and equal to the total of all cancer patients combined^2^. While advances in retinal disease diagnostics have progressed rapidly, specific treatments for AMD directed at underlying genetic or metabolic defects have progressed slowly due to limited understanding of disease pathways and cell types involved in the initiation of AMD. The retina lines the inner surface of the eye and neurally connects to the brain via the optic nerve (Fig. 1a). Photoreceptors and their support cells form a vertically organized, tightly integrated physiologic unit (Fig. 1b). AMD is a disease of this unit, with secondary effects including gliosis, cell death and synaptic circuity corruption in inner retina^3–5^. Given the complexity of the retinal cell structure, there is an urgent need to identify cells contributing to the spectrum of AMD pathology.

**Figure 1.**
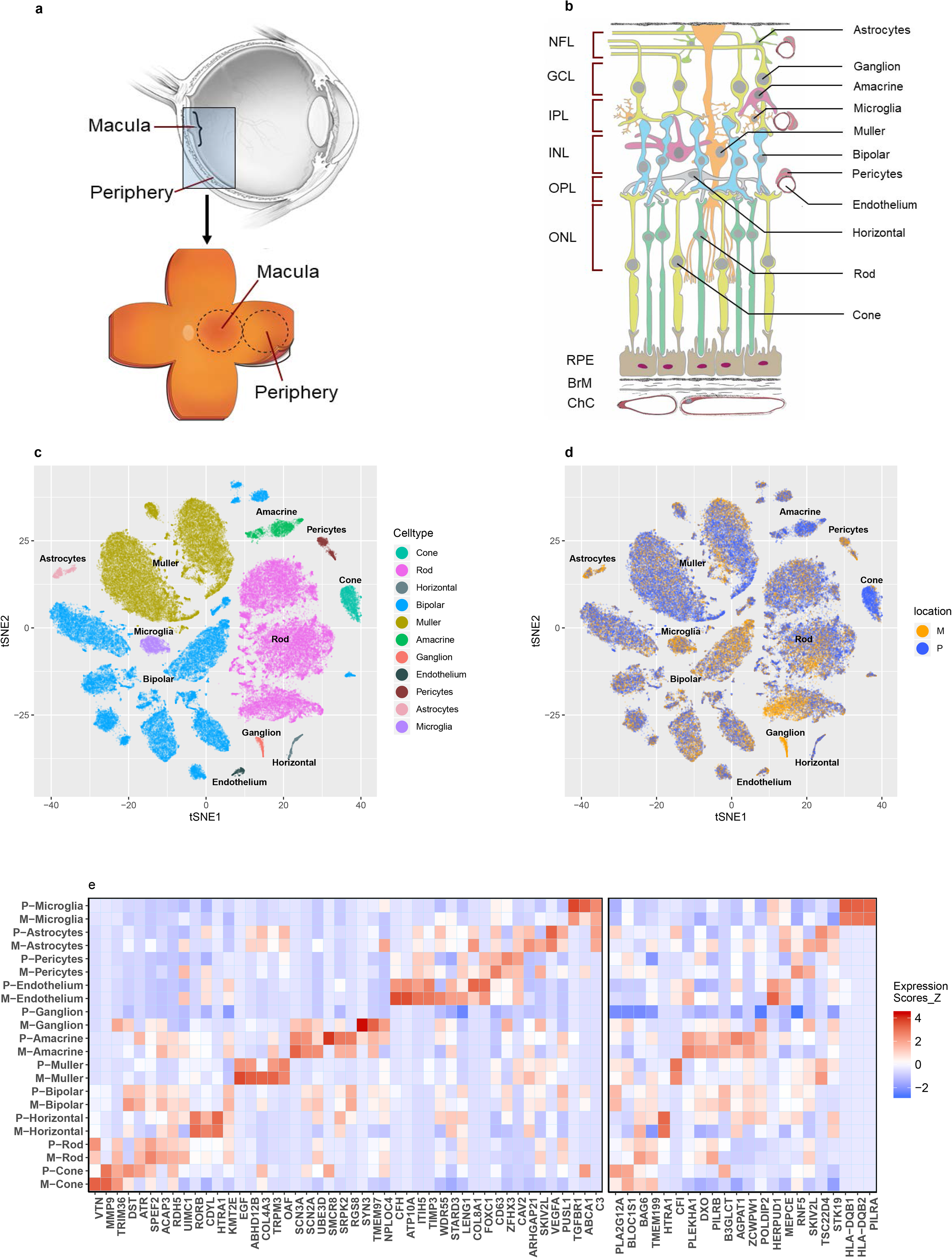
Summary of single-cell analysis from human retina. (a) Schematic cross-section of human eye (top) showing the retina lining the interior surface. The macula contains the fovea and is responsible for sharp vision. The periphery is responsible for detecting light and motion. Schematic of dissected tissue (bottom) shows retina adjoined to support tissues, flattened with relaxing cuts. Areas 8 mm in diameter were excised for RNA sequencing. (b) Layers of human retina and supporting tissues showing 11 assayed cell types. Five neuronal classes are photoreceptors, bipolar cells, ganglion cells, horizontal and amacrine cells. Cone photoreceptors are sensitive to color and bright light. Rod photoreceptors are sensitive to low light. Ganglion cells transmit information to the brain. Horizontal cells and amacrine cells modulate signal from photoreceptors and bipolar cells, respectively. Müller glia span the retina and are involved in neurotransmission, fluid balance, and wound repair. Also depicted are microglia (with phagocytic and immune activity), astrocytes (regulation of metabolism and blood brain barrier, synaptogenesis, neurotransmission), vascular endothelium (vascular tone and blood flow; coagulation and fibrinolysis; immune response, inflammation and angiogenesis) and pericytes (integrity of endothelial cells, trans-regulation of vascular tone, stem cells). The retinal layers include: NFL, nerve fiber layer; GCL, ganglion cell layer; IPL, inner plexiform layer; INL, inner nuclear layer; OPL, outer plexiform layer; ONL, outer nuclear layer; RPE, retinal pigment epithelium; BrM, Bruch’s membrane; ChC, choriocapillaris. The last three are shown for completeness and were not assayed. (c) Visualization of single-cell clusters using t-SNE. Cells are colored by cell types. (d) Visualization of single-cell clusters using t-SNE. Cells are colored by region. Cells from macular and peripheral retina were randomly mixed, suggesting the absence of batch effect. (e) Heatmap showing expression levels of AMD risk genes by cell type. Color in the heatmap represents expression intensity with red signifying higher expression in units of z-score. Left panel: AMD associated genes identified by loss-or gain-of-function mutations or by GWAS^6^. Right panel: target genes based on TWAS analysis listed^10^.

Recent technologic breakthroughs in scRNA-seq make it possible to measure gene expression in single cells, resolve cell types, characterize the signature of gene expression across cells, and improve understanding of cellular function in health and disease^6–9^. We performed scRNA-seq on macula and peripheral retina from two postmortem normal eyes. In total, we obtained 36,959 macular and 55,426 peripheral cells from the retina. Unsupervised deep learning based clustering identified 11 broadly defined cell types (Fig. 1c). Although some neuronal cell types, such as bipolar, amacrine, and ganglion cells can be further subdivided, we omitted further sub-clustering and maintained major cell types.

We assessed the cell-type specificity of 75 AMD GWAS^6^ and transcriptome-wide association study (TWAS)^10^ risk genes (**Fig. 1e, Methods and Supplementary Fig. 4**). Of 75 AMD risk genes, 23 showed cell type-specific expression either in macula or peripheral retina in the scRNA-eq data (**Methods, Supplementary Data 2**). For example, *MMP9* is specifically expressed in cones, and *PILRA* and *HLA-DQB1* are preferentially expressed in microglia. Further, 41 AMD risk genes have significant differential expression (DE) for retinal location (adjusted *P* <0.05) across 11 cell types.

To investigate the impact of AMD on cell types, we sequenced total RNA from macula and peripheral regions of 15 postmortem retinas that included normal, early and advanced AMD stages. Retinas were phenotyped by *ex vivo* fundus imaging and fellow-eye histology at The University of Alabama at Birmingham (UAB). Analysis identified 9,772 and 1,214 differentially expressed genes (DEGs) in macula and periphery for normal vs late AMD comparison (**Supplementary Data 3**). Interestingly, the DEG analysis between normal and early AMD found 169 DEGs in periphery and 21 genes in macula. We expected to see more DEGs in the macula than periphery and suspect that the larger sample size and higher sequencing depth in the peripheral retina samples increased the power. Among DEGs identified in macula from either comparison, 1202 (12.3%) are cell type specific, and 183 (14.6%) DEGs identified in periphery show cell type specificity (**Methods, Supplementary Fig. 5b and Supplementary Note 6**). Interestingly, we also found 17 DEGs for macula that may associate with AMD progression, as indicated by their increased fold change from early AMD to late AMD when compared to normal (**Supplementary Data 3**). Three of the AMD progression-associated DEGs show cell type-specificity, including RAB41 (Cone), ZMYND19 (Rod) and COL4A3 (Müller glia).

It is known that AMD also has an impact on cell type composition of the retina, particularly in macula^11,12^. To characterize such changes in cell type composition, we estimated cell type proportions for each bulk RNA-seq sample by cell type deconvolution analysis using MuSiC in which the scRNA-seq data was used as a reference^13^. First, we considered a large bulk RNA-seq dataset generated by the EyeGEx study^10^, which includes 453 RNA-seq samples generated from peripheral retina in postmortem human donors. This dataset includes samples at different AMD stages based on the Minnesota Grading System (MGS) (MGS1: 105; MGS2: 175; MGS3: 112; MGS4: 61). For the normal eyes (MGS1), our deconvolution analysis revealed a noticeable proportion of rod photoreceptors (mean proportion=0.58) and Müller glia (mean proportion=0.14), but relatively small proportions (0.01<mean proportion<0.1) of ganglion cells, cones, amacrine, bipolar, astrocytes and horizontal cells (Fig 2a). As AMD progresses (from MGS 2 to MGS 4) rods decrease. In contrast, the proportion of astrocytes increase, possibly reflecting an immune response^14^ of the peripheral retina to AMD.

**Figure 2.**
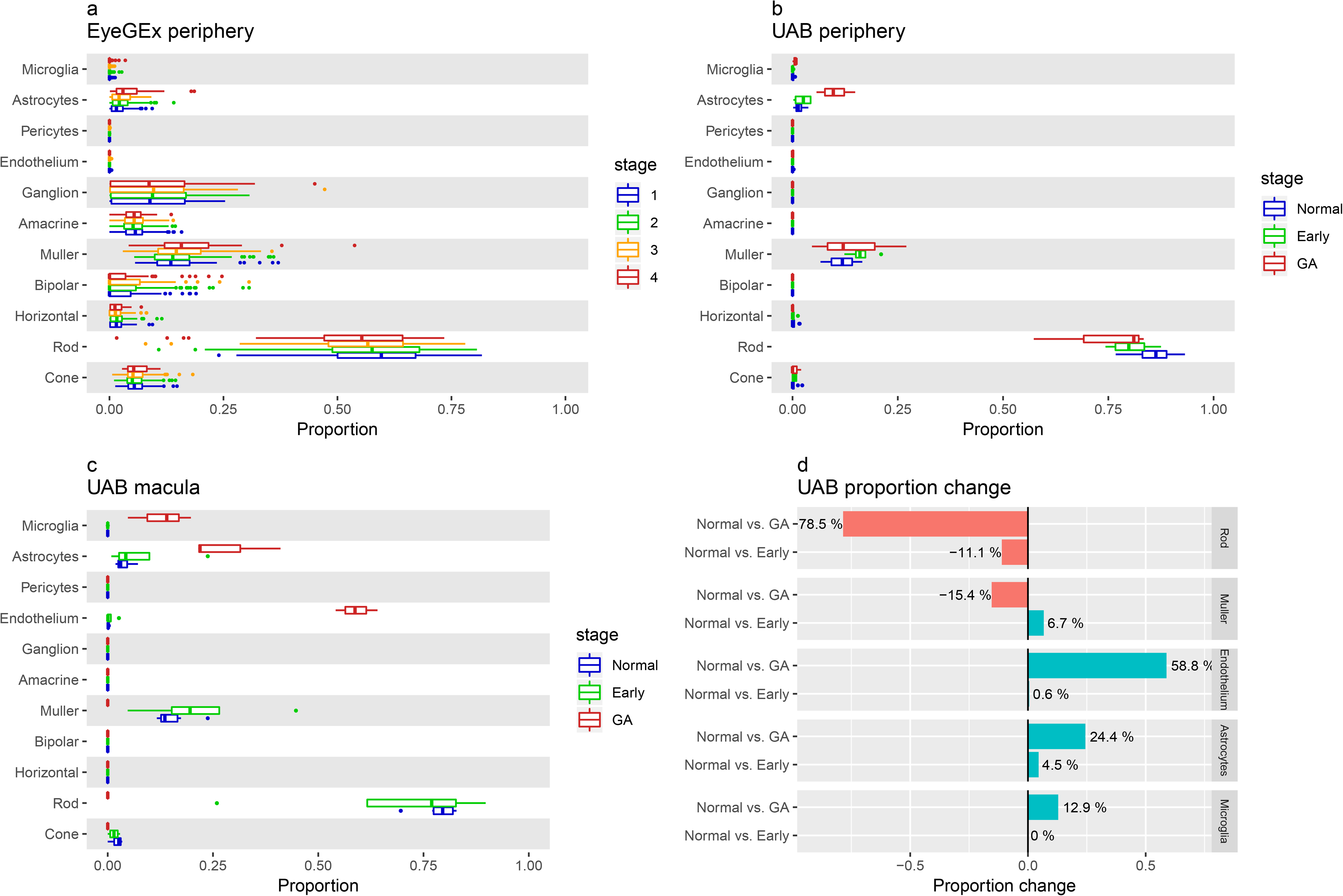
Cell type deconvolution analysis from bulk RNA-seq data. Cell type proportions for each bulk RNA-seq sample were estimated using MuSiC with the scRNA-seq data as reference. (a) Estimated cell type proportions for the EyeGEx peripheral retina bulk RNA-samples with four stages of AMD (MGS1: 105; MGS2: 175; MGS3: 112; MGS4: 61). (b) Estimated cell type proportions for the UAB peripheral retina bulk RNA-seq samples (normal: 8; early AMD: 4; late AMD: 3). (c) Estimated cell type proportions for the UAB macular retina bulk RNA-seq samples (normal: 6; early AMD: 4; late AMD: 3). Note the similarity in (a) and (b) with respect to cell proportion increase in astrocytes and decrease in rods in peripheral retina as AMD progresses. Larger differences are noted in both cell types in macula along with additional increases in Mϋller glia, microglia and vascular endothelium as AMD progresses. (d) Cell type proportion changes in the UAB macula retina samples for highlighted cell types.

The EyeGEx dataset is based is restricted to peripheral retina. Since AMD preferentially affects macula, we performed cell type deconvolution in our sample set which includes both macula and peripheral retina. In our peripheral retina samples, we found a decrease in rods and increase in astrocytes as AMD progresses, consistent with the EyeGEx data (Fig 2b). In macula, rods show a slightly decrease from normal to early AMD and a sharp decline from early to advanced AMD (Fig. 2c). Endothelium, astrocytes and microglia proportions increased in the macula when progressing from normal to advanced AMD. Rods are barely detectable in the macula of advanced AMD (Fig. 2d and Supplementary Fig. 6), in agreement with histological reports^11,15^.

As bulk RNA-seq measures the average expression of genes (sum of cell type-specific gene expression weighted by cell type proportions), DEGs from bulk RNA-seq can result from changes in cell type-specific gene expression, as well as cell type composition. To determine if DE in the bulk RNA-seq samples were due to cell type-specific DE and not change in cell type composition, we developed a calibration-based method to detect cell type-specific DEGs (ctDEGs) by calibrating bulk level gene expression using cell type-specific marker genes from the scRNA-seq data (**Methods**). Applying this method to the EyeGEx peripheral retina data, we detected AMD associated ctDEGs for each of the 11 major cell types. Across all cell types we identified 109 ctDEGs for MGS2 vs. MGS1, 201 ctDEGs for MGS4 vs. MGS1 (**Supplementary Data 4**), Fifty-one ctDEGs share the same cell type specificity for all comparisons. Only three ctDEGs were detected in astrocytes when comparing MGS3 vs. MGS1, possibly due to phenotype heterogeneity of the MGS3 samples. The largest set of cell type-specific DEGs were identified for microglia, 32 genes detected in MGS2 vs. MGS1 comparison, 82 genes detected for the MGS4 vs. MGS1 comparison (Fig. 3a), while 21 genes are in common and sharing the same directions of DE effect between two comparisons. Noticeably, for these 21 microglia-specific DEGs shared by two comparisons, the degree of fold change is generally higher in MGS4 vs. MGS1 than in MGS2 vs. MGS1, especially for *FCGBP* and *HLA-DME* (Fig. 3b and Supplementary Fig. 8). The increased expression of these genes in MGS4 reflects microglia-specific AMD response with disease progression. We observed a similar tendency for astrocytes- and endothelium-specific DEGs (**Supplementary Fig. 8**). We further performed Gene Ontology (GO) enrichment analysis on cell type-specific DEGs identified between MGS4 vs. MGS1. The results revealed distinct functional enrichment for up- and down- regulated genes (**Supplementary Data 5**); for example, microglia-specific up-regulated genes are enriched for immune response (adjusted *P* = 5.34×10^−17^), antigen processing and presentation of peptide antigen (adjusted *P* = 8.95×10^−14^) and innate immune response (adjusted *P* = 2.11×10^−12^), while down-regulated genes are enriched for nuclear-transcribed mRNA catabolic process and nonsense-mediated decay (adjusted *P* = 5.27×10^−14^), and establishment of protein localization to endoplasmic reticulum (adjusted *P* = 7.65×10^−13^).

**Figure 3.**
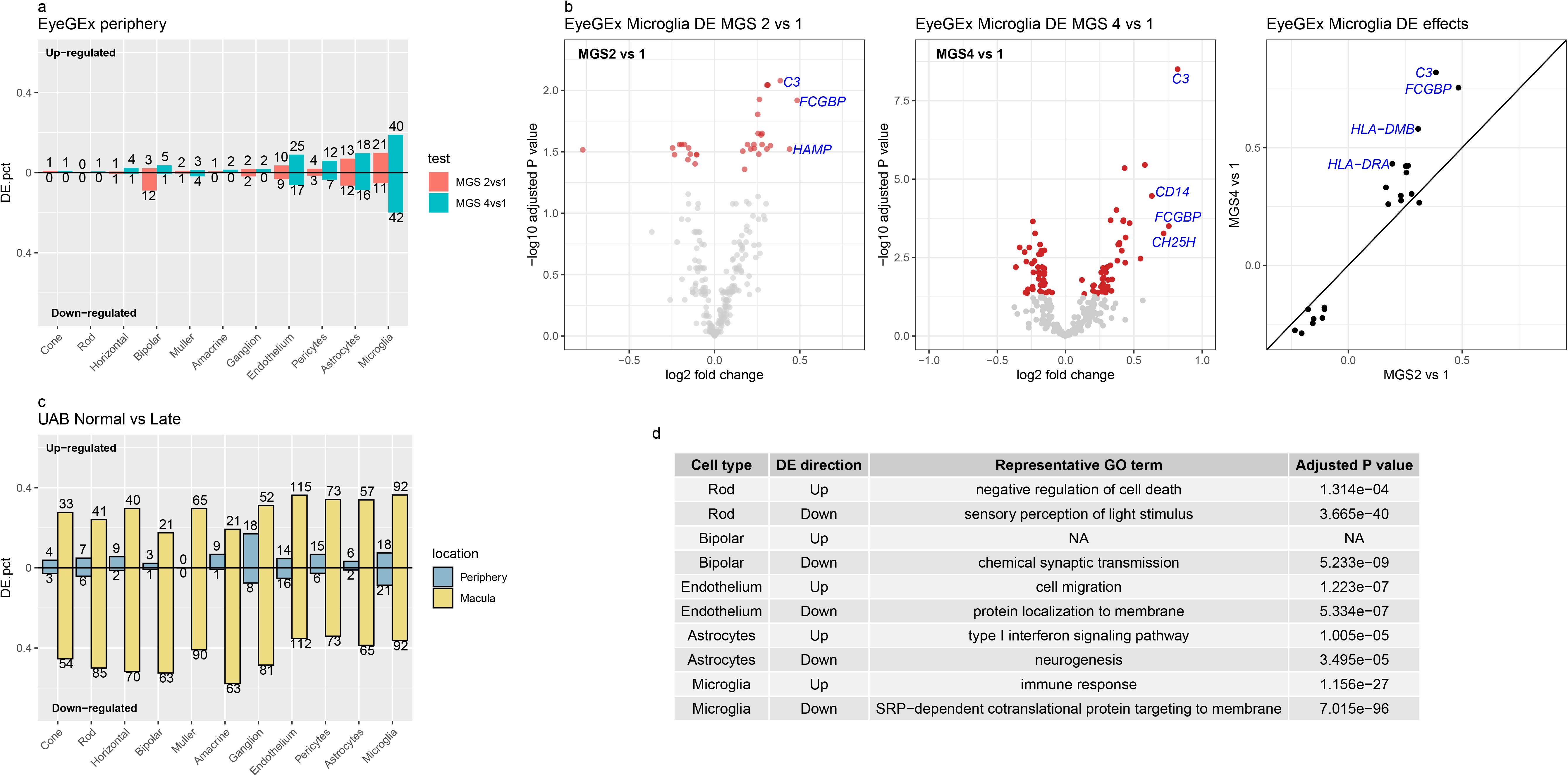
Cell type-specific differential expression analysis. (a) Proportions of up- and down-regulated ctDEGs detected identified in the EyeGEx peripheral retina data. Colors show different test conditions used in the DE analysis; red: MGS2 vs. MGS1, green: MGS4 vs. MGS1. (b) Volcano plots and effect size comparison of microglia-specific DEGs identified using EyeGEx peripheral retina data. Left/Mid: Volcano plots of microglia-specific DEGs identified from two different tests. Significant ctDEGs are highlighted using red color and ctDEGs with large effect size are annotated. Right: effect size comparison of microglia specific DEGs. X-axis: effect size of ctDEGs identified between MGS2 vs. MGS1; y-axis: effect size of ctDEGs identified between MGS4 vs. MGS1. ctDEGs with increased effect size are annotated. (c) Proportions of up- and down-regulated ctDEGs between normal vs. late AMD identified in UAB retina data. Colors show different retina regions in the DE analysis; red: periphery, green: macula. (d) Representative GO terms of up- and down-regulated ctDEGs genes identified between normal vs. late AMD using UAB data. The complete table of GO analysis result can be found in **Supplementary Data 5 and 7**.

To investigate the cell type-specific impact of AMD on macular retina, we applied our calibration-based ctDEG detection method to the UAB data. Due to the limited sample size and moderate alterations in expression pattern in early AMD, 7 and 13 ctDEGs were identified in macula and periphery, respectively, when comparing normal and early AMD (**Supplementary Data 6**). In contrast, a larger number of ctDEGs were detected when comparing normal and late AMD, with 169 ctDEGs found in periphery and 1,458 ctDEGs in macula (Fig. 3c). Among the 160 ctDEGs detected in periphery, a considerable number of them are microglia-(n=39) and endothelium-specific (n=30), which replicate the results in the EyeGEx periphery data. For macular retina, larger numbers (proportions of analyzed genes) of ctDEGs were identified for amacrine, bipolar, rod, cone and ganglion cells (Fig. 3c). This reflects the focus of AMD degeneration in the macula. Comparing DE of ctDEGs identified in the two regions we noticed a coordinated gene expression change between macula and periphery for immune related cell types (astrocytes and microglia) and photoreceptors (cones and rods) which indicate a greater impact on these cell types in the macula (**Supplementary Note 7 and Supplementary Fig 9**). Further, GO enrichment analysis on cell type-specific DEGs revealed that up- and down-regulated genes had distinct biological functions (**Supplementary data 7**). For example, microglia-specific up- and down-regulated DEGs between normal vs. late AMD identified in macula show a similar functional enrichment pattern as in the EyeGEx peripheral retina data (**Supplementary Data 5 and 7**). In particular, 41 rod specific up-regulated genes are enriched for negative regulation of cell death (adjusted *P* = 1.13×10^−2^) and negative regulation of apoptotic process (adjusted *P* = 3.56×10^−2^), whereas the 85 down-regulated genes are enriched for visual perception (adjusted *P* = 2.25×10^−40^), sensory perception of light stimulus (adjusted *P* = 3.65×10^−40^), and detection of light stimulus (adjusted *P* = 2.11×10^−33^) (Fig. 3d). These results reveal an impact on transcriptional profiles in different cell types that are common between macula and periphery, as well as transcriptional changes that are specific to each region.

In summary, we constructed a high-resolution human retina cell atlas with a particular focus on a comparison of regional differences. Our results linked GWAS genes for AMD with cell type-specific gene expression and enabled the use of GWAS data to inform the genetic architecture of AMD. We further leveraged scRNA-seq and bulk RNA-seq data, and our integrative analysis revealed both cell type-specific composition as well as gene expression changes associated with AMD progression. Our ongoing studies will aim to increase AMD sample size and add data from the RPE and choroid. Findings will overall provide novel insights into cell type-specific functions that can power precision therapeutic targeting of AMD.

## Acknowledgements

This work was supported by the following grants: NIH R01GM108600 (M.L.), R01GM125301 (M.L.), P30 EY003039 (UAB), R01EY030192 (M.L. and D.S.) and Macula Vision Research Foundation (D.S.). We thank the UAB comprehensive flow cytometry core for their services, and the Arnold and Mabel Beckman Initiative for Macular Research supported the collection of UAB eyes (C.A.C., D.S.).

## Author contributions

This study was conceived of and led by M.L. and D.S.. Y.L., R.Z., N.D., K.W., and Z.M. performed data analysis with input from M.L., D.S., and C.A.C.. C.S., S.L., and P.G. generated the human retina scRNA-seq data. N.P., Y.L. and M.L. created R Shiny apps for data visualization. Y.L. and M.L. wrote the paper with feedback from D.S., R.Z., C.A.C., K.W., Z.M., C.S., P.G..

## Methods

### Study subjects, scRNA-seq and bulk RNA-seq for the UAB data

The scRNA-seq data were generated from macular and peripheral retina taken from two healthy adult donors using the 10X Genomics Chromium^™^ system. The bulk RNA-data were generated from 13 macula samples (6 normal, 4 early AMD, and 3 late AMD) and 15 periphery samples (8 normal, 4 early AMD, and 3 late AMD) taken from the retina of 15 adult donors. All donor eyes were collected and characterized within 6 hours postmortem for presence of AMD and other pathology by author C.A.C. and a consulting medical retina specialist. Detailed sample preprocessing, donor characteristics, scRNA-seq and bulk RNA-seq data generation can be found in **Supplementary Note 1**.

### EyeGEx bulk RNA-seq data

The Eye Genotype Expression (EyeGEx) study was designed to explore genetic landscape and post-GWAS interpretation of multifactorial ocular traits^10^. This study generated bulk RNA-seq data of 523 peripheral retinal samples from postmortem human donors. We obtained the EyeGEx bulk RNA-seq data from the Gene Expression Omnibus (accession number GSE115828). This dataset includes gene expression measures for 523 samples and 58,051 genes. 453 of the samples with AMD phenotype information (MGS1: 105; MGS2: 175; MGS3: 112; MGS4: 61) were included in the analysis^16^. Genes that were expressed in less than 20% of the samples were eliminated, resulting in 14,709 genes in downstream analyses.

### scRNA-seq data clustering and cell type identification

To identify cell types in the scRNA-seq data, we clustered cells into distinct cell types using DESC, a deep learning based clustering algorithm that is robust to batch effect^17^. To prepare the data for DESC clustering, the original gene count matrix obtained from CellRanger was normalized in which the UMI count for each gene in each cell was divided by the total number of UMIs in the cell. The normalized UMI count data were then multiplied by 10,000 and transformed to a natural log scale. We further standardize the log-transformed expression value for each gene by calculating a Z-score across cells within each batch. Lastly, 2,000 highly variable genes selected using *filter_genes_dispersion* function from the Scanpy package^18^ were used as input for DESC clustering. In DESC analysis, we used a 2-layer autoencoder with 64 nodes on the first layer and 32 nodes on the second layer. The DESC clustering was performed using a grid of resolutions, and resolution = 0.4 was selected because it yields high maximum cluster assignment probability for most of the cells. DESC initially identified 18 cell clusters and 16 of them that contain more than 50 cells were kept for downstream analyses. We annotated these 16 cell clusters with cell type labels by examining expression patterns of known retina cell type markers (**Supplementary Data 8**). We further performed pairwise differential expression analysis among cell clusters, and cell clusters with the same cell type annotation and very few differentially expressed genes were merged (**Supplementary Note 2 and Supplementary Fig. 2**). This procedure resulted in 11 major neuronal cell types, including cone photoreceptors, rod photoreceptors, bipolar cells, horizontal cells, amacrine cells, and ganglion cells; support cells (microglia, Mϋller glia, astrocytes), and vascular cells (endothelium, pericytes).

We are aware of that some of the cell types we identified, such as cone, rod and ganglion, are commonly called cell classes,^19^ since each of them includes multiple (sub) types of cells with different expression patterns. However, to simplify the analysis of neural and non-neural cells, we use **cell type** to signify both **cell types** and **cell classes** in our data.

### t-SNE visualization for single-cell clustering

To visualize cell type clusters from the scRNA-seq data, we generated a two-dimensional non-linear embedding of the cells using t-distributed Stochastic Neighbor Embedding (t-SNE)^20^. The low denominational representation of the original data from DESC were used as input. The algorithm was implemented using the mTSNE function from python package MulticoreTSNE^21^. We set perplexity = 50 and learning rate = 500 and used the default values for all other parameters.

### Cell type- and region-specific expression of AMD risk genes

We obtained AMD risk genes from previous studies, which include 51 AMD associated GWAS genes from Peng et al. 2019^6^ and 26 target genes identified from TWAS analysis by Ratnapriya et al. 2019^10^. A gene that meets the following criteria was included for downstream analysis: 1) expressed in at least 1% of the cells; 2) expressed in at least 10 cells for at least one cell type in the scRNA-seq data. In total, 46 AMD associated genes and 22 TWAS target genes met these criteria. For these 68 genes, we tested whether they have significant high expression levels in a particular retina region and cell type(s). To test the region specificity, for each cell type, we tested whether these genes are differentially expressed between two retina regions. The analysis was conducted using the *FindAllMarkers* function in the R Seurat package^22^ with the Wilcoxon test. Benjamini-Hochberg (BH)^23^ adjusted p-value < 0.05 was used as threshold. To examine the cell type specificity, for each region, we compared AMD risk gene list to the identified cell type-specific genes and counted the overlap (**Supplementary Note 3 and Supplementary Data 1**).

To visualize cell type and region-specific expression, for each AMD risk gene, we calculated the mean expression across cells for each of the 11 major cell types for macular and peripheral retina separately. If a gene is expressed in less than 1% or 15 cells in a particular cell type, the mean expression of this gene in this cell type will be set as 0. Genes that have 0 mean expression across all cell types will be removed from further analysis. To make cell type-wise mean expressions comparable across genes, we calculated z-score of cell type mean expressions for each gene, and visualized the z-scores using heatmap (Fig. 1e).

### DEG detection in bulk RNA-seq data

Differential expression analysis for bulk RNA-seq data was performed using DEseq2 (v1.22.2)^24^. For the EyeGEx data, the filtered RSEM count matrix (14,709 genes by 453 samples) was used as input. Differential expression analysis was performed between normal vs. AMD samples defined by three different MGS levels. Also, sex was included as a covariate in the analysis. For the UAB data, we detected DEGs for macula and periphery separately. Genes that were expressed in less than 20% of the samples were eliminated, resulting in 19,313 genes in downstream analyses. The filtered read count matrices (19,313 genes by 13 samples for macula; 19,313 genes by 15 samples for periphery) were used as input. For each retina region, we detected DEGs between normal vs. early and normal vs. late AMD. All parameters for DESeq2 were set as default. We used BH adjusted p-value < 0.05 as significance threshold to correct for multiple testing. The significant DEGs are reported in **Supplementary Data 3**.

Further, we also examined whether these DEGs are cell type specific. For each retina region, we counted the overlap between identified DEGs and cell type-specific genes. Then we reported the proportions of cell type specific ones in AMD associated DEGs.

### Cell type deconvolution in bulk RNA-seq data

We performed cell type deconvolution analysis for both the EyeGEx and UAB bulk RNA-seq data using the UAB scRNA-seq data as the reference. For the scRNA-seq data, we only kept genes that were expressed in at least 5% of cells and more than 10 cells in at least one cell type. Cell type deconvolution analysis was conducted using MuSiC^13^ by setting eps = 0.0001, iter.max = 1,000 and default values for all other parameters. Also, we used the collection of 1,701 cell type-specific marker genes as reference genes in the deconvolution (**Supplementary Note 3 and Supplementary Data 1**).

### Detection of cell type-specific DEGs using calibrated gene expression

Our analysis shows AMD may have specific impact on particular cell types. We are interested in detecting differential expression between normal and AMD eyes for different cell types separately. However, the bulk RNA-seq data with both normal and AMD subjects lack cell type level information. To bypass such limitation, we developed a procedure to detect cell type-specific DEGs using bulk RNA-seq data calibrated by cell type proportion, which can be obtained from scRNA-seq data.

Consider a scenario in which we aim to calculate fold change of gene expression between two conditions for a particular cell type. Let *Y_gi_* denote the bulk RNA-seq expression for gene *g* in sample *i*. *Y_gi_* is a weighted sum of cell type level gene expression,

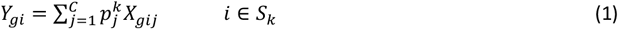

where 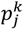 is the proportion of cell type *j* (*j* = 1, 2,…,*C*) under condition *k* (*k* = 1,2), *X_gij_* is expression level of gene *g* in sample *i* for cell type *j*, and *S_k_* is the set of samples under condition *k*. Here we assume that if gene *g* is cell type *c* specific, it is only expressed in that cell type so that

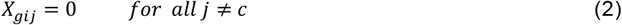

Combine (1) and (2), then for genes that are cell type *c* specific, we have:

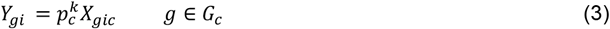

where *G_c_* is the set of genes that are cell type *c* specific. Let *Z_g_* denote the fold change of gene *g* between two conditions in cell type *c*. Then

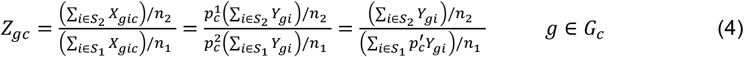

where 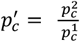 is the proportion change of cell type *c* between the two conditions, *n_k_* (*k* = 1,2) is the number of samples in condition *k*. Thus, for cell type *c* specific gene *g*, the cell type level fold change *Z_gc_* can be calculated using bulk level expression *Y_gi_* calibrated by 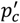, which is the proportion change of cell type *c*.

For each of the 11 cell types, we aim to identify ctDEGs. Firstly, we calibrate bulk expression levels for identified cell type-specific markers (**Supplementary Note 3 and Supplementary Data 1**) according to (4), and then performed differential expression analysis for these genes using DEseq2^24^. All parameters in DESeq2 were set at default and genes with Benjamini-Hochberg (BH)^23^ adjusted p-value < 0.05 was declared to be significant. The detected cell type specific DEGs are reported in **Supplementary Data 4 and 6**.

### Alternative way to calculate cell type proportion change

Although we are able to estimate proportion change by averaging the between-condition difference of cell type proportion the from the deconvolution results, the proportion change obtained this way is subject to sample variation, prone to outliers, and it may result in larger number of false positives in the detected cell type specific DEGs. Therefore, we propose an alternative way to estimate cell type proportion change which can increase the robustness of cell type specific DEGs detection.

We assume that for a given cell type, only few cell type specific markers are differentially expressed between conditions for the cell type, and the average fold change across genes specific to the cell type is 1, that is,

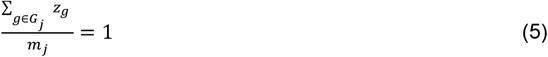

where *m_j_* is the number of cell type specific genes for cell type *j*. Combine (4) and (5) we have:

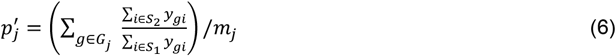

Thus, we are able to calculate between-condition proportion change 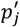 for cell type *j* directly using bulk level expression for cell type specific markers. Under our assumption, this method is a more direct way to estimate cell type proportion change between conditions. By avoiding the sample variation and complexity introduced in the deconvolution analysis, the calculation is more robust. The method was applied in the procedure of cell type specific DEGs detection (**Supplementary Fig.7**).

### GO enrichment analysis for ctDEGs

We preformed GO-enrichment analysis using ToppGene Suite (https://toppgene.cchmc.org/)^25^ for up- and down- regulated genes specific to each cell type. The analysis was perfomed only if there are at least 10 ctEGDs in the list. We used Bonferroni corrected P-value < 0.05 as the threshold for the significant GO-terms. The results are reported in **Supplementary data 5 and 7**. Representative GO terms for rod, bipolar, endothelium, astrocytes and microglia are shown in Fig. 3d.

