## Supplementery note & figures for "Integrative single-cell and bulk RNA-seq analysis in human retina identified cell type-specific composition and gene expression changes for age-related macular degeneration"

[1]Department of Biostatistics, Epidemiology and Informatics, University of Pennsylvania Perelman School of Medicine, Philadelphia, PA 19104, USA; [2]Department of Chemistry and Biochemistry, The University of the Sciences in Philadelphia, Philadelphia, PA 19104, USA; [3]Dept of Ophthalmology and Human Genetics, University of Pennsylvania Perelman School of Medicine, Philadelphia, PA 19104, USA; [4]Department of Psychology, University of Alabama at Birmingham, Birmingham, AL 35294, USA; [5]Department of Biochemistry and Molecular Genetics, University of Alabama at Birmingham, Birmingham, AL 35294, USA; [6]Department of Computer and Information Science, University of Pennsylvania, PA 19104, USA; [7]Department of Ophthalmology and Visual Sciences, University of Alabama at Birmingham, AL 35294, USA.

[*] Equal contribution

### Supplementary Notes

Supplementary Note 1. scRNA-seq sample processing and data generation
Supplementary Note 2. scRNA-seq data clustering and cell type assignment
Supplementary Note 3. Identification of cell type-specific marker genes
Supplementary Note 4. Bulk RNA-seq sample processing and data generation
Supplementary Note 5. Comparison of UAB and EyeGEx bulk RNA-seq data
Supplementary Note 6. AMD associated DEGs are tend to be cell type-specific
Supplementary Note 7. Comparison of ctDEGs effect across disease stages and regions

### Supplementary Figures

Supplementary Fig. 1. Overview and quality control of scRNA-seq data
Supplementary Fig. 2. Similarity in expression pattern across cell clusters
Supplementary Fig. 3. Identified cell types from scRNA-seq data
Supplementary Fig. 4. Expression of AMD risk genes across cell types and retina regions

Supplementary Fig. 5. Similarity of bulk RNA-seq data across datesets and conditions

Supplementary Fig. 6. Cell type proportion changes across AMD stages

Supplementary Fig. 7. DE fold change for cell type markers in bulk RNA-seq data level

Supplementary Fig. 8. Comparison of ctDEGs effect between AMD stages

Supplementary Fig. 9. Comparison of ctDEGs effect between retina regions

Supplementary Fig. 10. Comparison of ct DEGs effect between periphery retina datasets.

**Supplementary Data**

Supplementary Data 1. Cell type-specific gene markers detected from the scRNA-seq data.

Supplementary Data 2. Single-cell level expression patterns of AMD risk genes.

Supplementary Data 3. Differential expression results for the UAB bulk RNA-seq data.

Supplementary Data 4. ctDEGs identified in the EyeGEx bulk RNA-seq data.

Supplementary Data 5. GO enrichment result for ctDEGs in the EyeGEx bulk RNA-seq data.

Supplementary Data 6. ctDEGs identified in the UAB bulk RNA-seq data.

Supplementary Data 7. GO enrichment result for ctDEGs in the UAB bulk RNA-seq data.

Supplementary Data 8. Known retina cell type markers.

### Supplementary Note 1. scRNA-seq sample processing and data generation

**Eye collection protocol.** Donor eyes from two Caucasian male donors, aged 78 and 90, were obtained from Advancing Sight Network (formerly Alabama Eye Bank) within 6 hours postmortem. Neither donor had a known history of retinal disease, head or ocular trauma, significant refractive error, neurological disease, diabetes, or uncontrolled hypertension.

**Dissection and dissociation of retina.** Following removal of the anterior chamber and vitreous, the eyecup was immersed in oxygenated Ames media. Relief cuts were made in the posterior eyecup to expose the retinal tissue and 8 mm punches were obtained from macular retina and temporal (peripheral) retina and carefully isolated from the combined retinal pigment epithelium and choroid tissue. The isolated neurosensory retina was dissociated with activated papain (Worthington Biochemical Corp.) as previously optimized to obtain a high percentage of viable retinal cells[1]. After dissociation, magnetic bead-based removal of dead cells (Miltenyi Biotech) was used to reach the optimum target for viability of 85-95% per sample. Viability was determined by FACS sort or by staining an aliquot of the dissociated cells with trypan blue, 0.4% (Sigma-Aldrich).

**Single cell transcriptome library preparation and sequencing.** Single cell transcriptome libraries were prepared by using 10xGenomics Single Cell 3' biased v2 kit according to the company's manual. The constructed single cell libraries were sequenced by HiSeq 2000 sequencer (Illumina, Inc., San Diego, CA, USA) with total reads per cell targeted for a minimum of 50,000.

**Preprocessing of scRNA-seq data.** For each sample replicate, we performed initial quality control using Cell Ranger (Version 2.1.0). For macular retina samples, we initially obtained a 33,694 genes by 48,800 cells count matrix. For peripheral retina samples, we obtained a 33,694 genes by 59,222 cells count matricx. Then, we further filtered the data using Seurat (version 2.3.4)[2]. A cell was retained in downstream analyses if it meets the following criteria: (1) More than 200 genes are detected; (2) The proportion of the transcript counts from mitochondrial encoded genes is less than 25%; (3) Total number of UMIs is between 500 and 10,000. This resulted in 33,694 genes across 92,386 cells (macula: 36,959; periphery: 55,426), which were used for analysis shown in **Supplementary Fig. 1**.

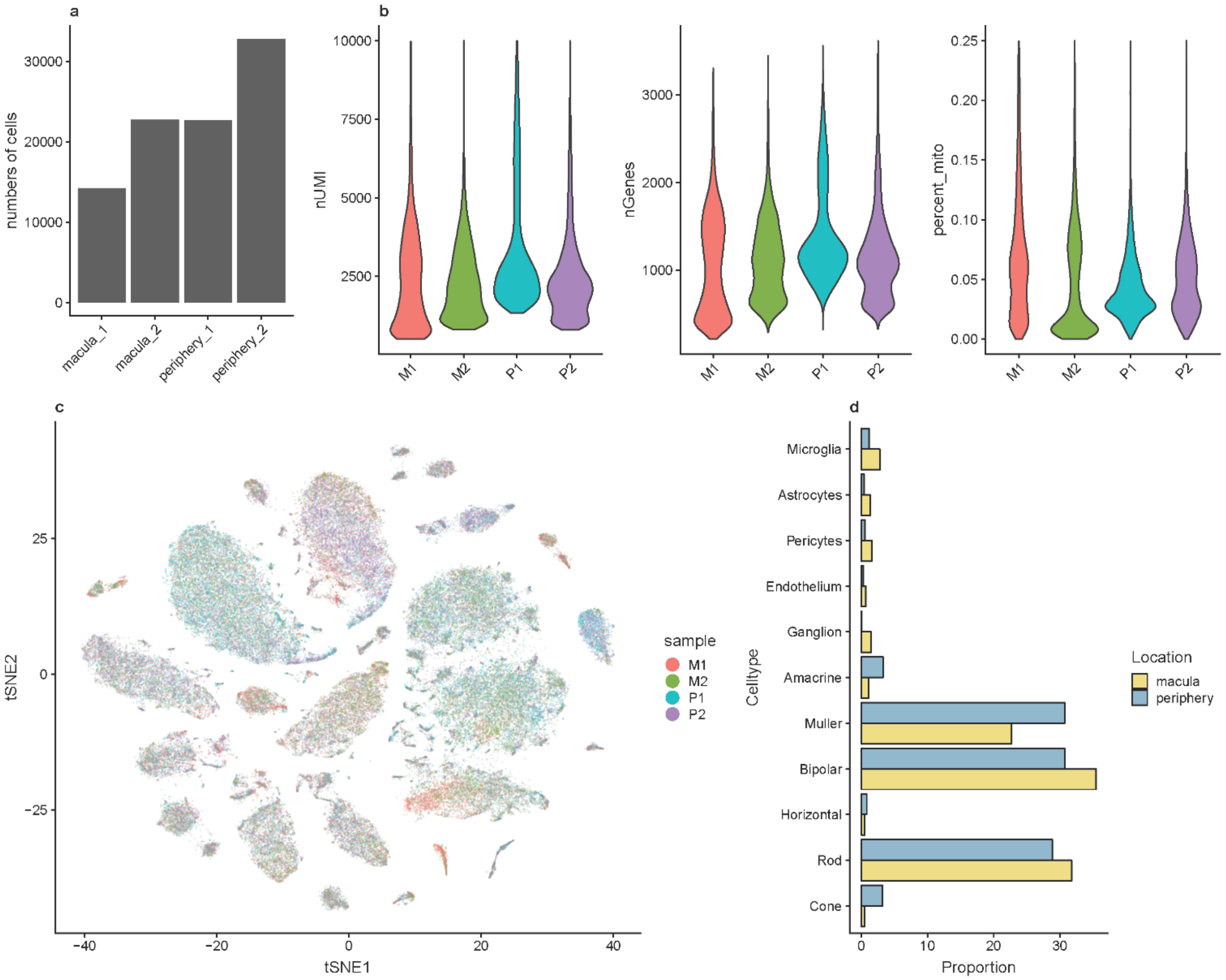


**Supplementary Fig. 1. Overview and quality control of scRNA-seq data.**

**(a)** Bar graph showing the number of cells for the four filtered scRNA-seq samples (two samples for macular retina and two samples for peripheral retina). **(b)** Violin plots showing the distribution of the number of UMIs per cell (left), number of genes per cell (middle) and percentage of mitochondrial genes per cell for the four scRNA-seq samples. **(c)** t-SNE projections of the scRNA-seq data. Color labels cells from different samples. The cells are randomly mixed, indicating batch effect was removed in clustering. **(d)** Bar plots showing proportions of identified cell types from the scRNA-seq data across the two retina regions.

**Supplementary Note 2. scRNA-seq data clustering and cell type assignment**

To identify cell types in the scRNA-seq data, we clustered cells into transcriptionally similar groups using DESC[3] (**Methods**). Initially, we obtained 18 cell clusters but decided to continue with 16 cell clusters because 2 cell clusters had less than 50 cells. We annotated these 16 cell clusters with cell type labels by examining expression patterns of known retina cell type markers (**Supplementary Data 8**). We identified six bipolar subtypes. To examine these 16 cell clusters further, we performed pairwise differential expression analysis among 16 cell clusters using *FindMarkers* from the Seurat package. We enabled the Wilcoxon test by specifying test.use = "wilcox" and all other parameters were set as default, and used adjusted $p\text{-value}<0.05$, $\text{fold-change}>2$ as threshold to determine significant DEGs between each pair of cell clusters. We detected a considerable number of DEGs between each pair of cell clusters, except the pairwise analysis among the 6 cell clusters labeled as bipolar (**Supplementary Fig. 2**). Therefore, we maintained these 6 clusters as bipolar cell subtypes. In total, we determined 11 major cell types. **Supplementary Fig. 3** shows the expression patterns of representative known cell type markers across the 11 identified cell types and the hieratical similarities of these cell types.

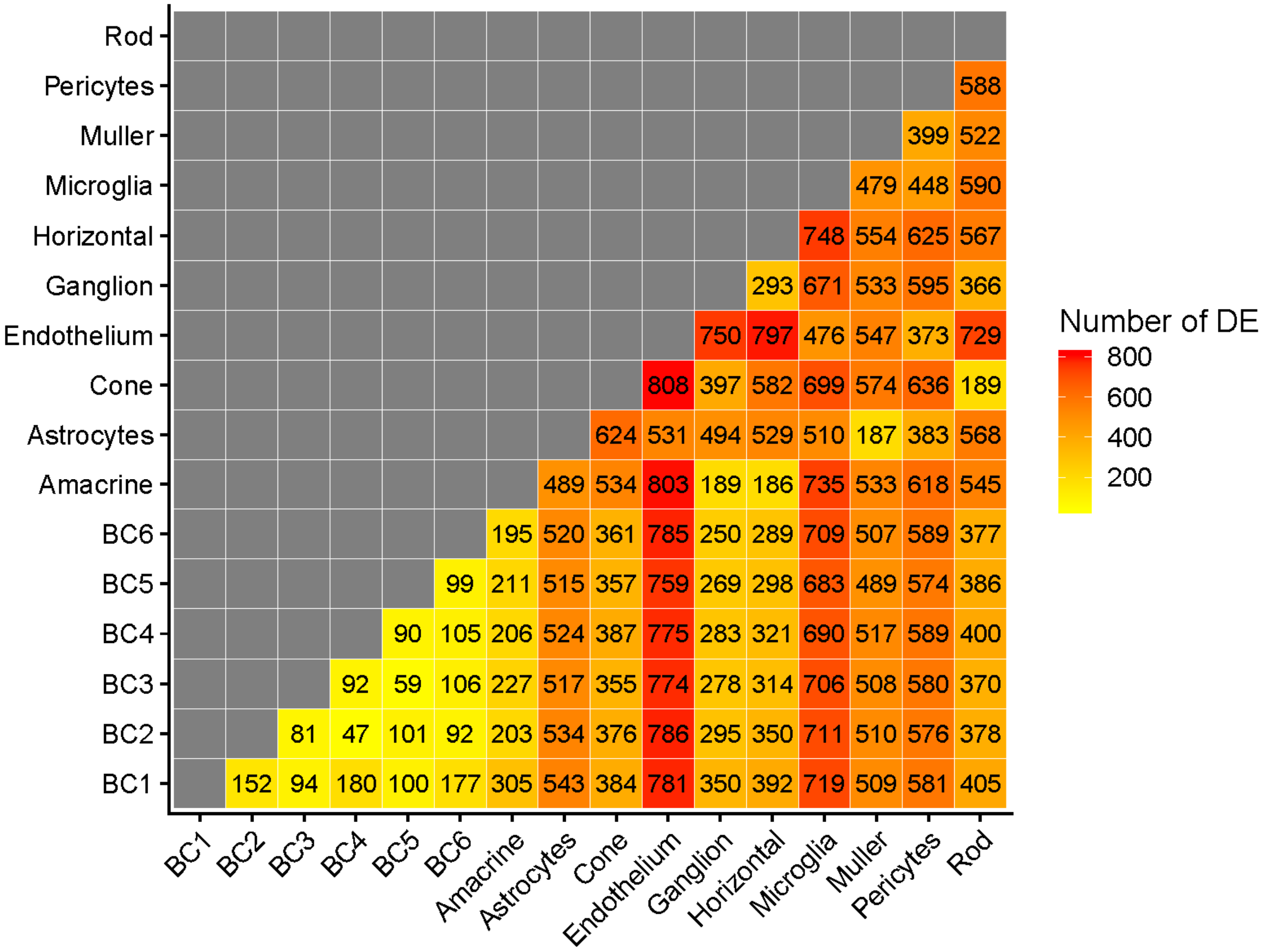


**Supplementary Fig. 2. Similarity in expression pattern across cell clusters.** The heatmap shows the number of significant DEGs detected between each pair of cell clusters. Sixteen cell clusters were identified using DESC and six of them were labeled as bipolar subtypes (BC1-BC6). The annotation in tiles show the exact number of significant DEGs for each pairwise differential expression analysis

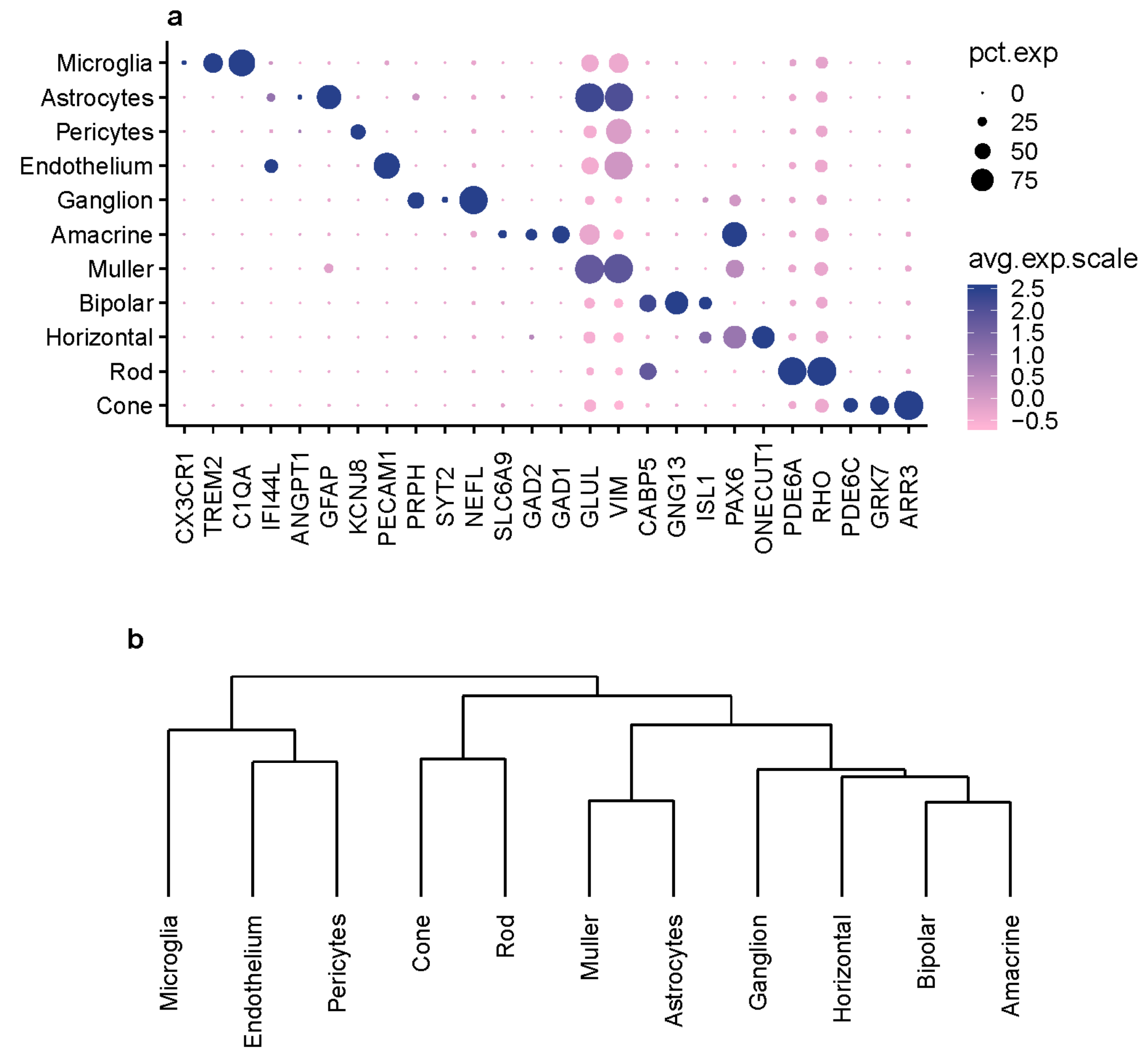


**Supplementary Fig. 3. Identified cell types from scRNA-seq data. (a)** Dot plots showing expression pattern of known gene markers across cell types identified (**Supplementary Data 8**). The dot plot was generated using *Dotplot* in R Seurat package. The size of the dots represents percentage of cells that expressed gene markers while color shows average expression levels of gene markers. **(b)** The dendrogram shows the hierarchy of identified cell types. Hierarchical clustering analysis was performed on the mean expression for all genes across cells within each of the cell types.

**Supplementary Note 3. Identification of cell type-specific marker genes**

To determine if a gene is preferentially expressed in a given cell type, we performed differential expression analysis to test whether a gene has a significantly higher expression in the given cell type than all other cell types. The analysis was implemented using the *FindMarkers* function in Seurat R package. We used the Wilcoxon test for the differential expression analysis by specifying test.use = "wilcox" and all other parameters were set as default. Then, the significant (adjusted P value<0.05, positive fold change>2) DEGs for each of the cell types were defined as cell type-specific genes. Note that it is possible that a gene is specific to multiple cell types, although such cases are rare.

We preformed the cell type-specific markers identification by combining data from two retina regions, as well as using macula and periphery data separately. Identified cell type-specific genes can be found in **Supplementary Data 1**.

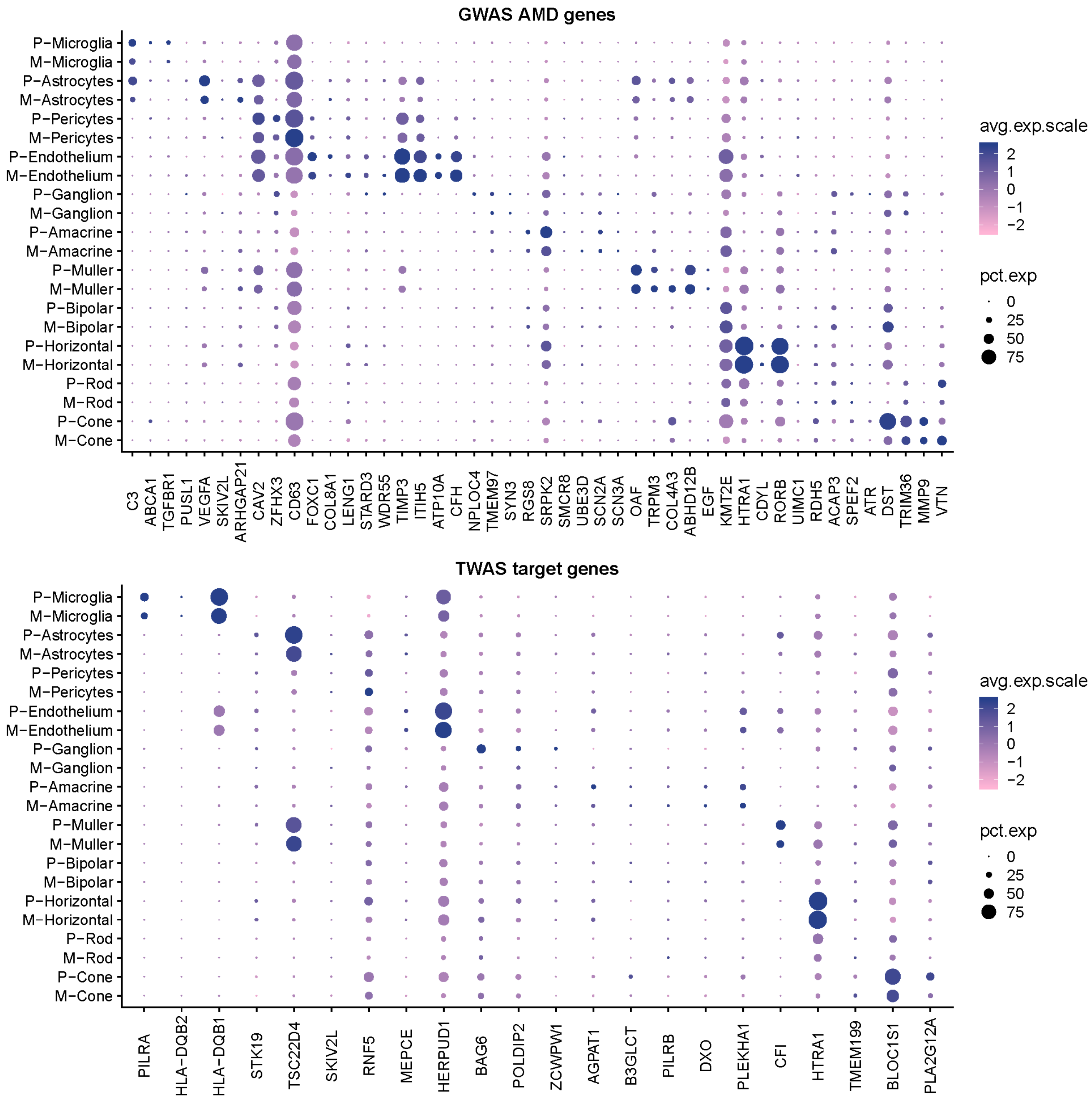


**Supplementary Fig. 4. Expression of AMD risk genes across cell types and retina regions.** Dot plots showing expression patterns of AMD risk genes (top: GWAS AMD gene, bottom: TWAS target genes) across cell types and retina regions. The dot plot was generated using *Dotplot* in R Seurat package. The size of the dots represents percentage of cells that expressed gene markers while color shows average expression levels of gene markers.

**Supplementary Note 4. UAB bulk tissue processing and data generation**

**Histopathological analysis of UAB samples.** The Institutional Review Board at UAB approved the use of human tissues in this study. This study utilized 8 pairs of eyes from non-diabetic Caucasian donors 69-95 yr of age (84.73 yr ± 5.53 yr; 8 males and 7 females) at a death-to-preservation interval of < 6 hr. Ocular health histories were not available. Eyes were opened by eye bank recovery personnel using an 18 mm diameter corneal trephine, followed by a snip to the iris to facilitate penetrate of preservatives into the fundus. Preservatives were RNAlater (Qiagen) for the Left Eye and 2% glutaraldehyde and 1% paraformaldehyde in 0.1M phosphate buffer for the Right Eye, both at 4°C. Left Eyes were shipped on wet ice via overnight courier to University of Pennsylvania where they were processed upon arrival.

Maculopathy status of Right Eyes was assessed at UAB by a 3-component protocol. Eyes underwent multimodal ex vivo imaging of excised 8-mm diameter macular punches using digital color photography and spectral domain optical coherence tomography volume scans (SD-OCT; Spectralis, Heidelberg Engineering) with a custom tissue holder (co-author JDM). They also underwent internal globe examination using a dissecting scope (Nikon SMZ-U) with oblique trans- and epi-illumination in consultation with an MD medical retina specialist (co-author JAK). Finally, eyes were submitted for histopathology using macula-wide high-resolution sections. Macular punches including retina, RPE, choroid, and sclera were then post-fixed in osmium tannic acid paraphenylenediamine to accentuate neutral lipid-rich lesions associated with age-related macular degeneration (AMD)[4,5]. Sections 0.8 µm in thickness through the rod-free foveola and the rod-rich perifovea at 2000 µm superior to the fovea were stained with toluidine blue, examined, and annotated.

The definition of AMD used in this study[6] was the presence of one large druse (>125 µm in diameter) in the macula or severe RPE changes in the setting of at least one druse or continuous basal linear deposit, with or without the presence of neovascularization and its sequelae. Eyes with geographic atrophy had at least one region 250 µm in diameter lacking a continuous RPE layer (but possibly containing 'dissociated' RPE[7]. Unremarkable eyes lacking characteristics of AMD or other chorioretinal disease as discernable in either histology or ex vivo imaging served as comparison eyes.

The use of fellow eyes optimally preserved for RNA-seq and for histopathology in this study is a limitation, because two eyes of one individual may be at different disease severity. We feel that this limitation is manageable, because it was recently found in a population-based cohort that was observed for 20 years, AMD severity in one eye was found to largely track AMD severity in the fellow eye at all stages of the disease[8]. This published study also found a <10% chance of lifetime occurrence of asymmetry >2 steps on the grading scale for color fundus photography.

**Dissection and dissociation of retina.** Macula and periphery were dissected from retina resulting in two samples per eye. The tissues were isolated at the macula and periphery using a circular 10-mm biopsy punch.

**UAB data library preparation and sequencing.** RNA for the eye tissues was extracted using the AllPrep DNA/RNA Mini Kit (Qiagen). Extracted RNA samples underwent quality control assessment using R6K Screen Tape on a 2200 Tape Station (Agilent, Santa Clara, CA, USA) and were quantified using Qubit 2.0 Fluorimeter from Life Technologies (Grand Island, NY). All RNA samples selected for sequencing had an RNA integrity number of ≥8. Strand-specific RNA library was prepared from 100 ng totalRNAusing the Encore Complete RNA-Seq library kit (Nugen Technologies, Inc., San Carlos, CA, USA) according to the manufacturer's protocol. RNA sequencing was performed at the Center for Applied Genomics at the Children's Hospital of Philadelphia per standard protocols. The prepared libraries were clustered and then sequenced using HiSeq 2000 sequencer (Illumina, Inc., San Diego, CA, USA) with four RNA-seq libraries per lane (2 × 101-bp paired-end reads).

**Preprocessing of bulk tissue data.**

The RNA-Seq data were aligned to the hg38 reference genome using GSNAP (version 2016-06-30) with known splice sites (SNP file build 147) taken into account. In order to eliminate mapping errors and reduce potential mapping ambiguity owing to homologous sequences, several filtering steps were applied. Specifically, we required the mapping quality score of ≥30 for each read, reads from the same pair were mapped to the same chromosome with expected orientations and the mapping distance between members of the read pair was 200,000 bp. Quality control analysis of the aligned data was performed using program RNA-SeQC. All subsequent analyses were based on filtered alignment files.

Per-gene counts were generated from the GSNAP alignments using the HTSeq-count program (version 0.6.0) using default 'union' mode and the HG38 reference genome.

**Supplementary Note 5. Comparison of bulk tissue and EyeGEx bulk RNA-seq data**

Here we examined similarity of overall expression pattern across datasets and retinal regions. We calculated mean read counts within each datasets/conditions for genes that existed in both datasets. Then pairwise Pearson correlation were calculated upon the $\log_2$ scaled mean read counts (**Supplementary Fig. 5**). In both UAB and EyeGEx data, we observed that the overall expression pattern are similar (Spearman correlation>0.95) across all conditions for the periphery region. Even across different datasets, the overall expression of retina periphery are similar (Pearson correlation>0.84). In contrast, the macular region shows a distinct expression pattern compared to the periphery. In the UAB datasets, such difference is highlighted when comparing macula and periphery within the same condition (Pearson correlation<0.67). Further, the expression of macula under late AMD seems to be an outlier, distanced from all others, suggesting a drastic change in expression.

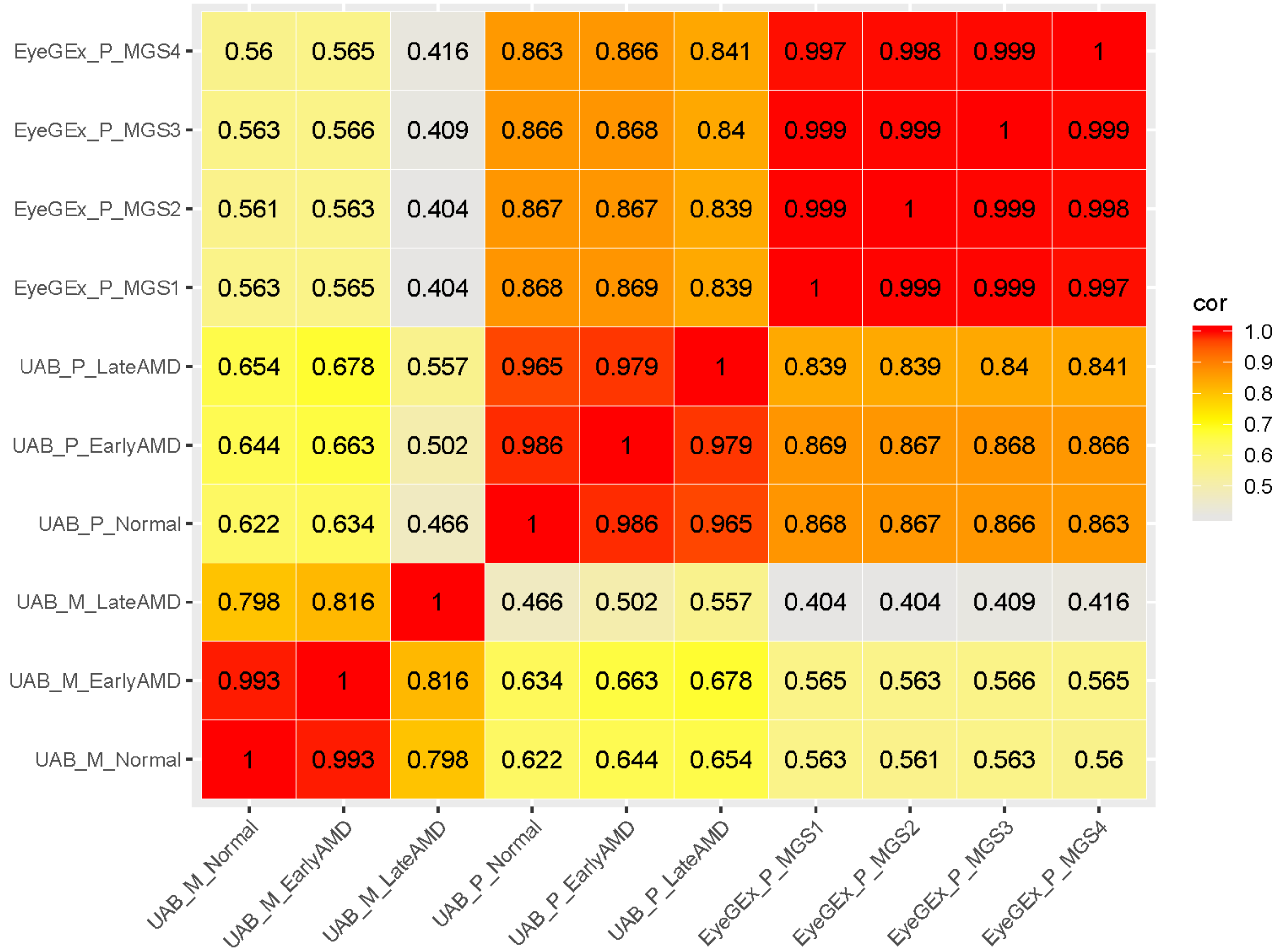


**Supplementary Fig. 5. Similarity of bulk RNA-seq data across datasets and conditions.** The heatmap shows the similarity (Pearson correlation) in overall expression pattern across UAB and EyeGEx datasets and conditions. Pearson correlation was calculated using log scales read counts between each pair of samples. Only genes that existed in both datasets were considered in the analysis.

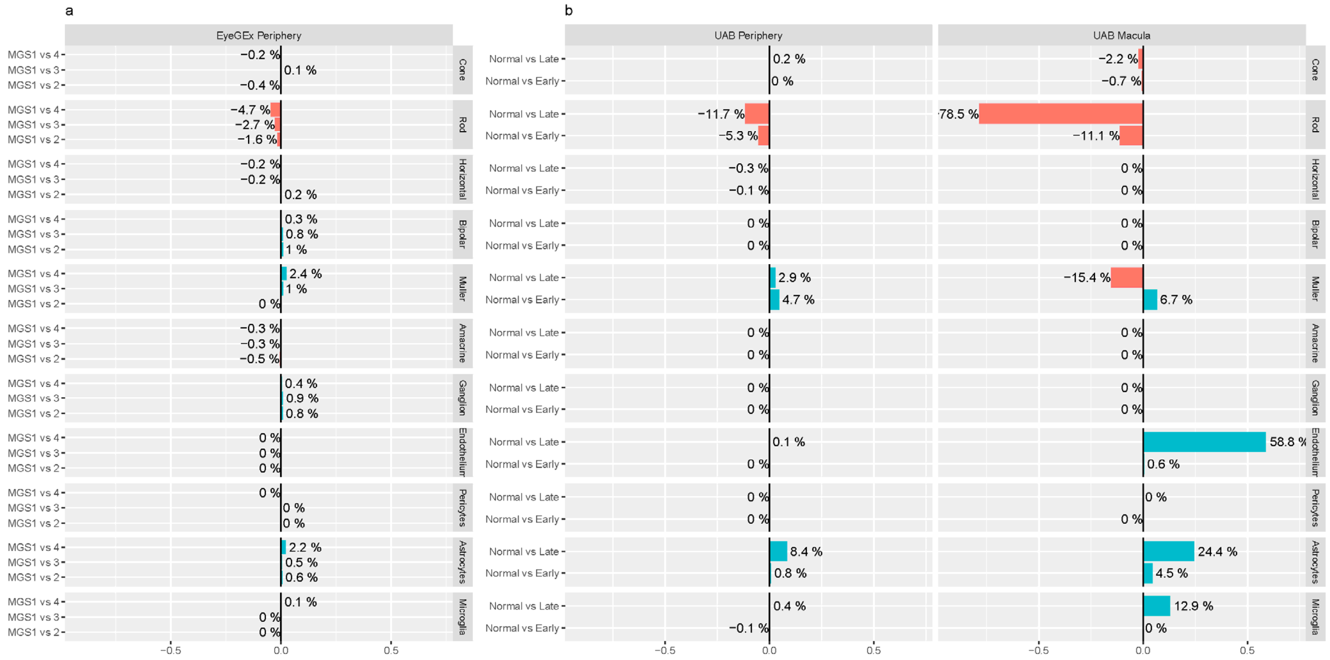


**Supplementary Fig. 6. Cell type proportion changes across AMD stages.** The bar graphs show proportion changes of cell type across different AMD stages. Color shows the direction of the changes (red: decreasing, green: increasing). **(a)** Changes in cell type proportions in EyeGEx data when comparing MGS 2, 3, and 4 to MGS 1. **(b)** Changes in cell type proportions in UAB data when comparing early and late AMD to normal.

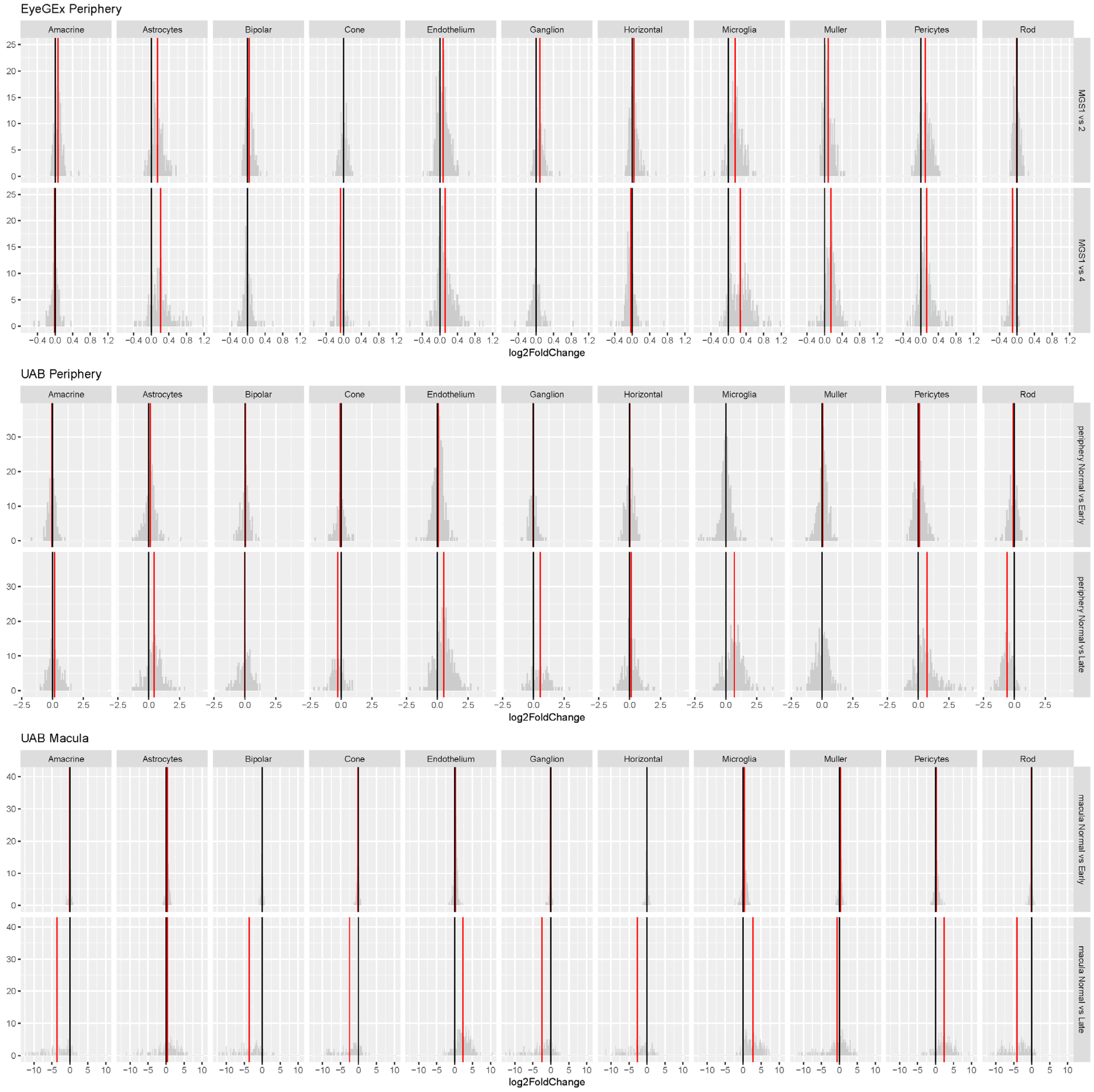


**Supplementary Fig. 7. DE fold change for cell type markers in bulk RNA-seq data level.** The histograms show the distribution of DE fold change for cell type markers in bulk RNA-seq data level (top: EyeGEx periphery, mid: UAB periphery, bottom: UAB macula). Red vertical line labels the mean of fold changes for each of the cell types while black vertical line labels $x = 0$ as comparison. For each of the cell type, genes that specifically expressed in the cell type were identified using scRNA-seq data. Then DE analysis were performed for these genes using bulk RNA-seq data. The calculated fold changes from the DE analysis were used to calibrate expression data in order to detected cell type-specific DEGs (**Methods**).

**Supplementary Note 6. Region- and cell type-specificity of AMD associated DEGs**

We counted the cell type specific genes (**Supplementary Note 3**) among identified DEGs using UAB data. For macula, 3 among 21 (14.2%) DEGs between AMD early and Normal, 1202 among 9772 (12.3%) DEGs between AMD late and Normal show cell type-specificity. For periphery, 10 among 169 (6.0%) DEGs between AMD early and Normal, 179 among 1214 (14.7%) DEGs between AMD late and Normal show cell type-specificity.

**Supplementary Note 7. Comparison of ctDEGs effect across disease stages, regions and datasets**

To investigate cell type-specific AMD impact, we identified genes that are differentially expressed in particular cell types **(Methods)**. These ctDEGs reflect the AMD response of different retinal cell types. It is possible that the level of such response are region-specific or relate to disease progression. To test our hypothesis, we examined linear relationship of log fold change between DE tests conducted for different AMD stages and retinal regions.

**Enhanced AMD response along with disease progression**. To compare the cell type specific AMD response between different disease stages, for each of the cell types, we examined linear relationship of log fold changes between two DE tests, MGS 2 vs. MGS1 and MGS 4 vs. MGS1, performed using EyeGEx data:

$$y_g^j = \beta^j x_g^j + \varepsilon_g^j \qquad g \in G_j$$

where $y_g^j$ is log fold changes of gene $g$ between MGS 4 vs. MGS1 at cell type $j$, while $x_g^j$ is the log fold change between MGS 2 vs. MGS1.The $\beta^j$ reflects the overall changes of DE effect size between tests for cell type $j$. In the analysis, p values were adjusted for the number of cell types using BH method. For each of the cell types, we reported estimated $\beta$ and its significance in the **Supplementary Fig. 8**. We identified significant $\beta$ (adjusted P value < 0.05) for amacrine, astrocytes, cone, endothelium, microglia, muller and pericytes. In cone ($\beta$=0.51), Müller ($\beta$=0.33) and pericytes ($\beta$=0.61), much larger DE fold changes were found between MGS 4 vs. MGS1 indicating an increased level of AMD response. While in endothelium and microglia, the fold change remain similar between DE tests for two AMD stage indicating a consistent AMD response for these two cell types.

**Enhanced AMD response in macula**. Similarly, to compare AMD response across retinal regions for, we compared fold changes between normal vs. late AMD calculated using UAB macula and periphery data:

$$y_g^j = \beta^j x_g^j + \varepsilon_g^j \qquad g \in G_j$$

where $y_g^j$ is log fold changes of gene $g$ between normal vs, late AMD at cell type $j$ calculated using macula data, while $x_g^j$ is the log fold change calculated using periphery data. The $\beta^j$ reflects the overall changes of DE effect size between retinal regions for cell type $j$. In the analysis, p values were adjusted for the number of cell types using BH method. For each of the cell types, we reported estimated $\beta$ and its significance in the **Supplementary Fig. 9**. Significant $\beta$ (adjusted P value < 0.05) were identified for astrocytes, cone, endothelium, microglia, pericytes and rod. We noticed a larger fold changes in neuron cell types including cone ($\beta$=0.04), horizontal ($\beta$=0.0.005) and rod ($\beta$=0.0.028).

**Consistent AMD response in periphery between datasets**. To compare AMD response in periphery retina across different datasets, we compared fold changes and between MGS 4 vs, MGS 1 calculated using EyeGEx periphery data, and between normal vs. late AMD calculated using UAB periphery:

$$y_g^j = \beta^j x_g^j + \varepsilon_g^j \qquad g \in G_j$$

where $y_g^j$ is log fold changes of gene $g$ between normal vs. late AMD at cell type $j$ calculated using UAB periphery data, while $x_g^j$ is the log fold change between MGS 4 vs. MGS 1 calculated using EyeGEx periphery data. In the analysis, p values were adjusted for the number of cell types using BH method. For each of the cell types, we reported estimated $\beta$ and its significance in the **Supplementary Fig. 10**. Significant (adjusted P value < 0.05) associate between fold changes estimated in different datasets were observed for all cell types that have large number of ctDEGs detected, including microglia, endothelium, astrocytes and pericytes. While such association are not significant for amacrine, Müller glia and rods which possibly due to the lower level-AMD response in periphery for these cell types. The result reveals the consistency between two periphery retina datasets.

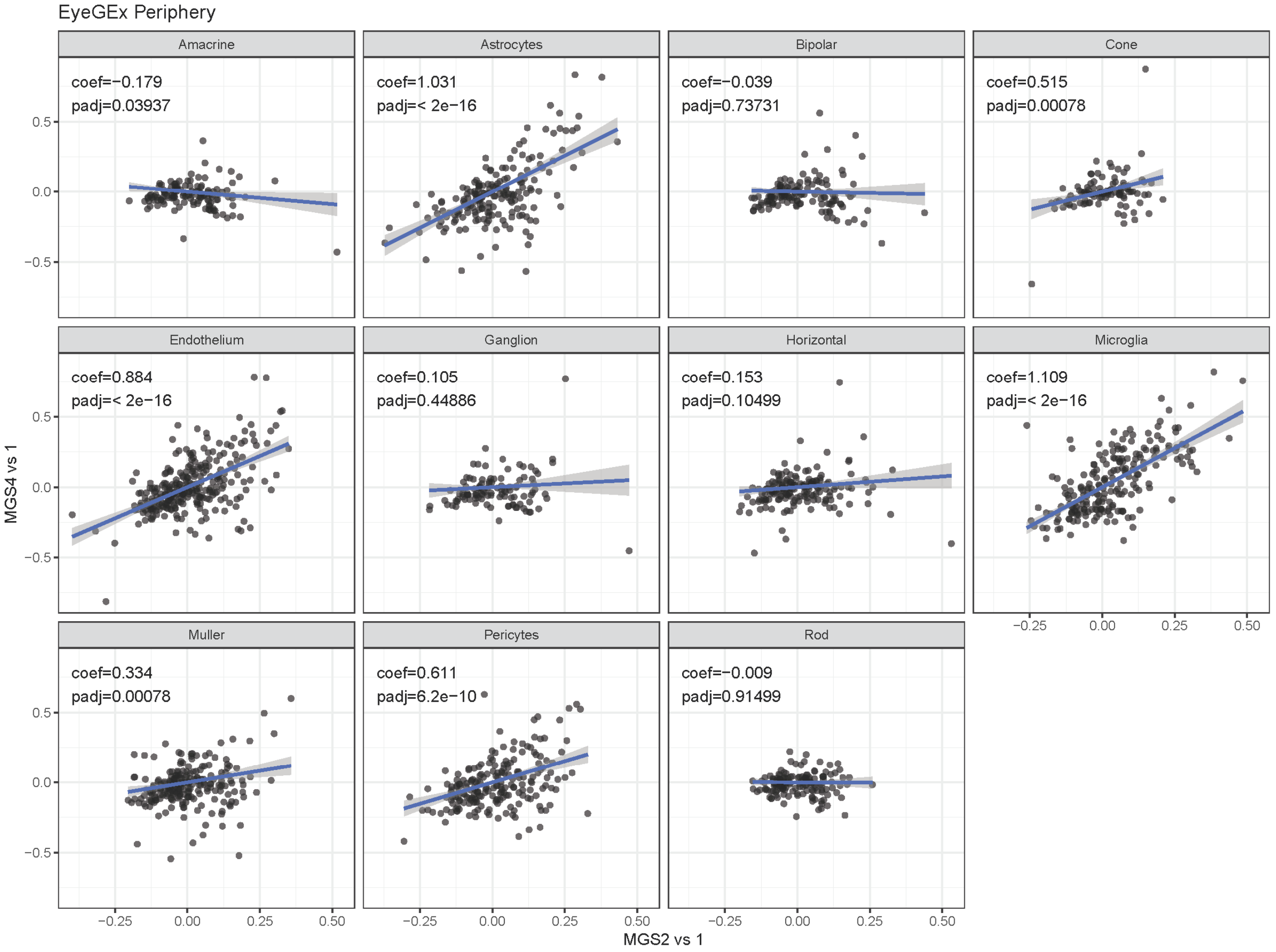


**Supplementary Fig. 8. Comparison of ctDEGs effect between AMD stages.** The scatter plots shows the comparison of effect size of ctDEGs identified from two test: MGS 2 vs.1 and MGS 4 vs. 1. For each of the cell types, regression was performed to investigate the possible linear relationship (**Supplementary Note 7**). Coefficients (coef) and adjusted p values (padj) for the linear regression were annotated on the plots.

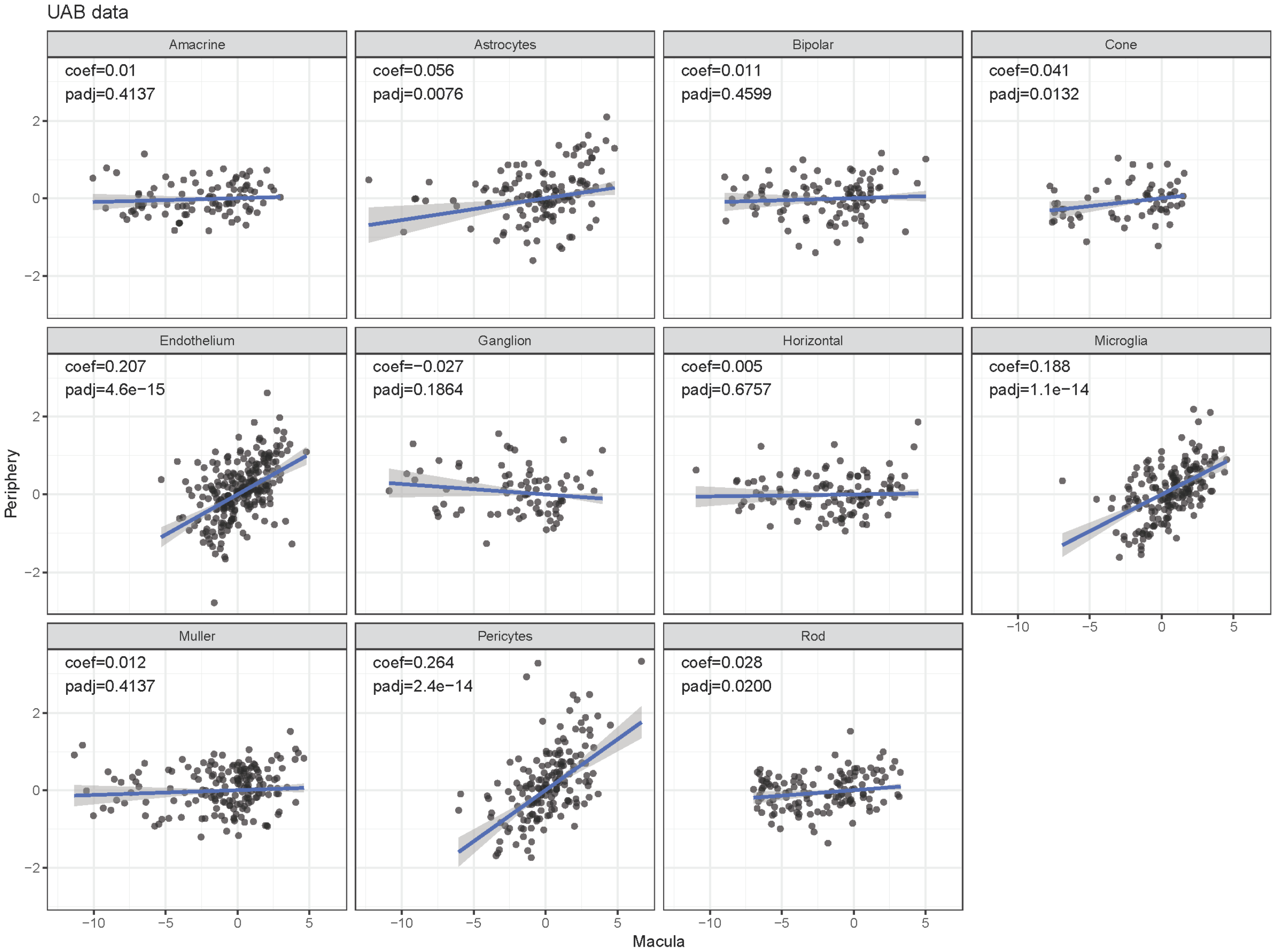


**Supplementary Fig. 9. Comparison of ctDEGs effect between retina regions.**

The scatter plots shows the comparison of effect size of ctDEGs identified from macula and periphery. For each of the cell types, regression was performed to investigate the possible linear relationship (**Supplementary Note 7**). Coefficients (coef) and adjusted p values (padj) for the linear regression were annotated on the plots.

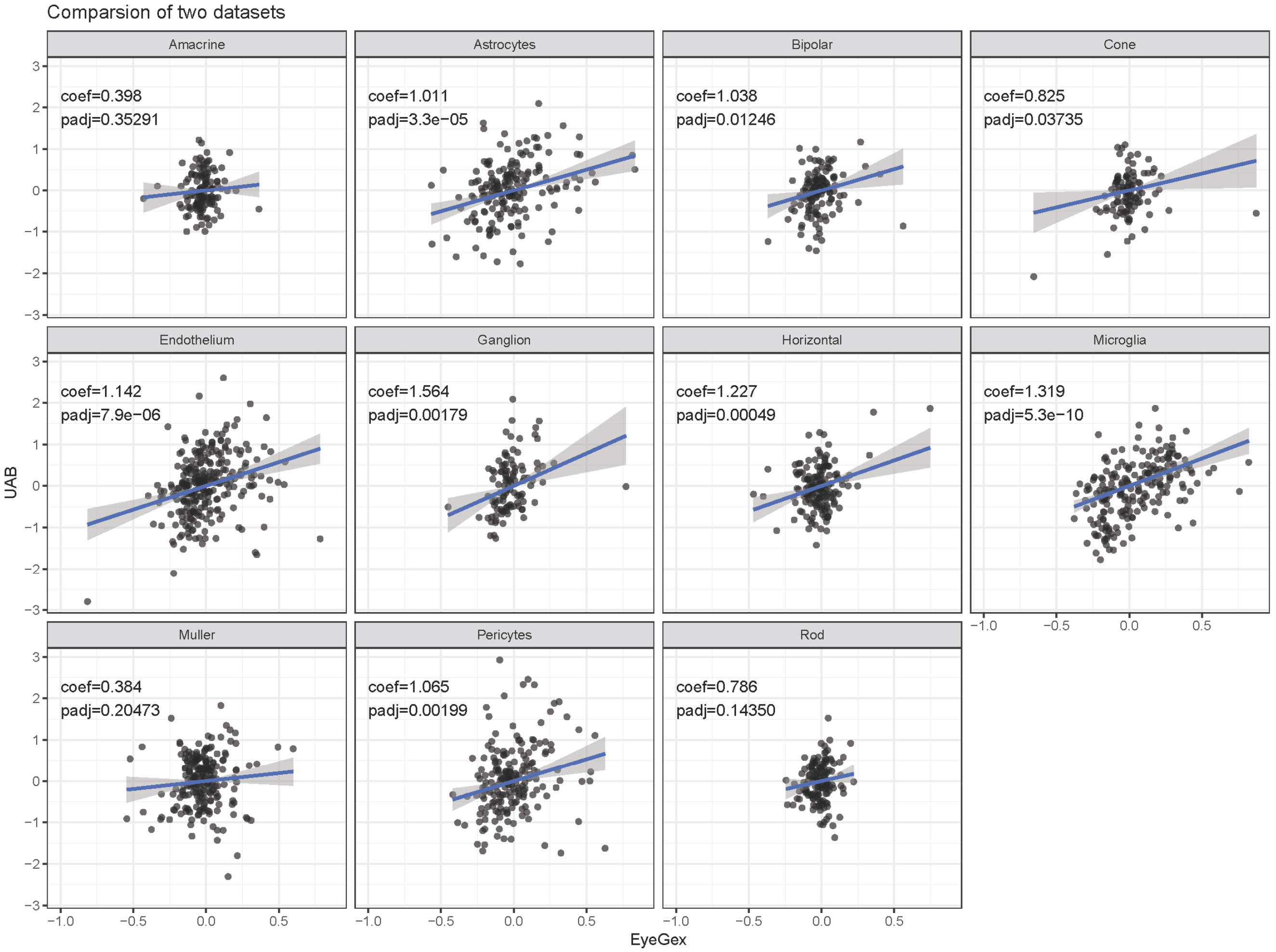


**Supplementary Fig. 10. Comparison of ctDEGs effect between periphery retina datasets.** The scatter plots shows the comparison of effect size of ctDEGs identified from different periphery retina datasets. For each of the cell types, regression was performed to investigate the possible associations (**Supplementary Note 7**). Coefficients (coef) and adjusted p values (padj) for the linear regression were annotated on the plots.

**Supplementary Data**

**Supplementary Data 1. Cell type-specific gene markers detected from the scRNA-seq data.**
a. Cell type-specific gene markers detected using combined data
b. Cell type-specific gene markers detected using macula data
c. Cell type-specific gene markers detected using periphery data

**Supplementary Data 2. Single-cell level expression patterns of AMD risk genes.**

**Supplementary Data 3. Differential expression results for the UAB bulk RNA-seq data.**

**Supplementary Data 4. ctDEGs identified in the EyeGEx bulk RNA-seq data.**
a. ctDEGs identified between MGS2 vs. MGS1
b. ctDEGs identified between MGS3 vs. MGS1
c. ctDEGs identified between MGS4 vs. MGS1

**Supplementary Data 5. GO enrichment result for ctDEGs in the EyeGEx bulk RNA-seq data.**

**Supplementary Data 6. ctDEGs identified in the UAB bulk RNA-seq data.**
a. ctDEGs identified between Early AMD vs. Normal in macula
b. ctDEGs identified between Late AMD vs. Normal in macula
c. ctDEGs identified between Early AMD vs. Normal in periphery
d. ctDEGs identified between Late AMD vs. Normal in periphery

**Supplementary Data 7. GO enrichment result for ctDEGs in the UAB bulk RNA-seq data.**

**Supplementary Data 8. Known retina cell type markers.**
